# Extracellular superoxide production is a widespread photoacclimation strategy in phytoplankton

**DOI:** 10.1101/2025.04.16.649151

**Authors:** Sydney Plummer, Susan Garcia, Julia M. Diaz

## Abstract

Phytoplankton control the habitability of Earth. These photosynthetic microorganisms serve as the base of marine food webs, produce approximately half of the planet’s oxygen, and regulate climate by sequestering carbon dioxide from the atmosphere. As global changes accelerate through the Anthropocene, phytoplankton communities face multiple stressors, including warming, shifting patterns in ocean circulation and structure, and associated perturbations in levels of light exposure. The health and functioning of the oceans depends on phytoplankton community responses to these stressors; however, the physiological processes involved in light stress are not fully understood. Here, we surveyed sixteen representative phytoplankton and show that most produce extracellular superoxide, an otherwise damaging reactive oxygen species, as a widespread strategy to acclimate to light stress. Indeed, all species regulated extracellular superoxide production as a function of light exposure, which was modeled with a modified photosynthesis-irradiance (PE) curve. Furthermore, the flavoenzyme inhibitor DPI quenched extracellular superoxide production and lead to declines in viability and photosynthetic health in thirteen out of sixteen species. The negative effect of DPI on photosynthetic health was stronger with increasing light, consistent with inhibition of a photoprotective process. Taken together, these results support the hypothesis that phytoplankton mitigate light stress through enzyme-mediated production of extracellular superoxide. These results imply that daytime rates of biological superoxide production in the marine environment are substantially underestimated by dark measurements. Furthermore, phytoplankton photoacclimation may alter superoxide production rates in future oceans impacted by changes in water column structure and light exposure.

## Introduction

Reactive oxygen species (ROS) form during the reduction of oxygen to water. These transient species are prevalent in aquatic environments like the oceans, where they arise via biological and non-biological pathways including photochemical production [1]. The reactive nature of ROS such as hydrogen peroxide (H_2_O_2_) and superoxide (O_2_^−^) has been linked to the transformation of important elements in aquatic systems. Indeed, superoxide and subsequent ROS decrease available oxygen [2] and regulate the speciation of harmful (e.g., Hg) [3] and nutrient metals (e.g., Fe, Cu) [4, 5]. ROS also contribute to carbon remineralization [1, 6, 7], especially through the action of hydroxyl radicals that can arise from superoxide [8]. In these ways, ROS influence ecosystem health and productivity.

All aerobic organisms make ROS. In fact, phytoplankton and bacteria are recognized as major sources of ROS in the marine environment [9] that can rival abiotic (i.e., photochemical) contributions [5, 10]. Biological ROS are often regarded as harmful byproducts of metabolism, yet ROS production can also benefit marine biota [8, 11–13]. Indeed, we recently reported that extracellular ROS (eROS) production is a beneficial photoacclimation strategy in the marine diatom *Thalassiosira oceanica*. We showed that *T. oceanica* uses a transmembrane protein to transfer electrons from cellular NADPH, a product of photosynthesis, to dissolved O_2_ on the outside of the cell, thereby making extracellular superoxide (eO_2_^−^). This process is dynamically regulated by *T. oceanica* as a function of light exposure and helps prevent cellular reductive stress under high irradiance [14].

It is unknown whether the eO_2_^−^-driven photoacclimation response observed in *T. oceanica* is widespread across phytoplankton taxa. Here, we surveyed a wide diversity of phytoplankton, including representatives of cyanobacteria and all major eukaryotic lineages, to determine how prevalent the photoacclimation role of eO_2_^−^ production may be. We show that all the tested phytoplankton regulate eO_2_^−^ production as a function of light exposure and that most of them use it as a beneficial strategy to photoacclimate. Because of this photoacclimation strategy, we suggest that future, climate-driven shifts in surface ocean habitats may alter biological rates of eO_2_^−^ production.

## Materials and Methods

### Phytoplankton cultivation

All phytoplankton were obtained from the National Center for Marine Algae and Microbiota (NCMA) at Bigelow Laboratory for Ocean Sciences, except for the following strains: *Karenia brevis* ARC5 from the Algal Resources Collection at the University of North Carolina Wilmington (www.algalresourcescollection.com), *Ostreococcus tauri* OTH95 from the Palenik lab (Scripps Institution of Oceanography), *Dunaliella* sp. 15-1a from the Bowman lab (Scripps Institution of Oceanography), *Phaeodactylum tricornutum* CCAP 1055/1 from the Allen lab (J. Craig Venter Institute), and *Prochlorococcus marinus* NATL2A and MIT9312 from the Chisholm lab (Massachusetts Institute of Technology). Phytoplankton were grown in autoclaved (121°C, 20 – 40 min) SN [15], L1 with the addition of silicic acid [16], f/2, f/2 with the addition of silicic acid [17], or Pro99 [18] media using 0.2 μm filtered natural seawater as a base (Supplementary Table 1). All experimental cultures were begun with stationary phase inoculum except for *P. marinus* strains, which were begun with exponential phase inoculum. Cultures were maintained in borosilicate culture tubes with caps at 18°C or 23°C under cool, white light (14:10 light dark cycle) (Supplementary Table 1). Growth was monitored by observing *in vivo* chlorophyll fluorescence using a handheld *Aqua*fluor® fluorometer (Turner Designs, San Jose, CA, USA) or cell abundance (cells mL^−1^) using a Guava® easyCyte flow cytometer (Luminex, Austin, TX, USA). Flow cytometry samples were analyzed by running live samples in a 96-well plate at a low flow rate of 0.24 μL s^−1^ for 3 minutes or until at least 1000 particles were counted. Instrument performance was validated daily using instrument specific beads. For analyses, gates of cell populations were created based on diagnostic red fluorescence and forward scatter signals of healthy, exponentially growing cells (exemplified by Supplementary Fig. 1). All experiments were conducted on exponentially growing cells, unless otherwise stated.

**Fig. 1.**
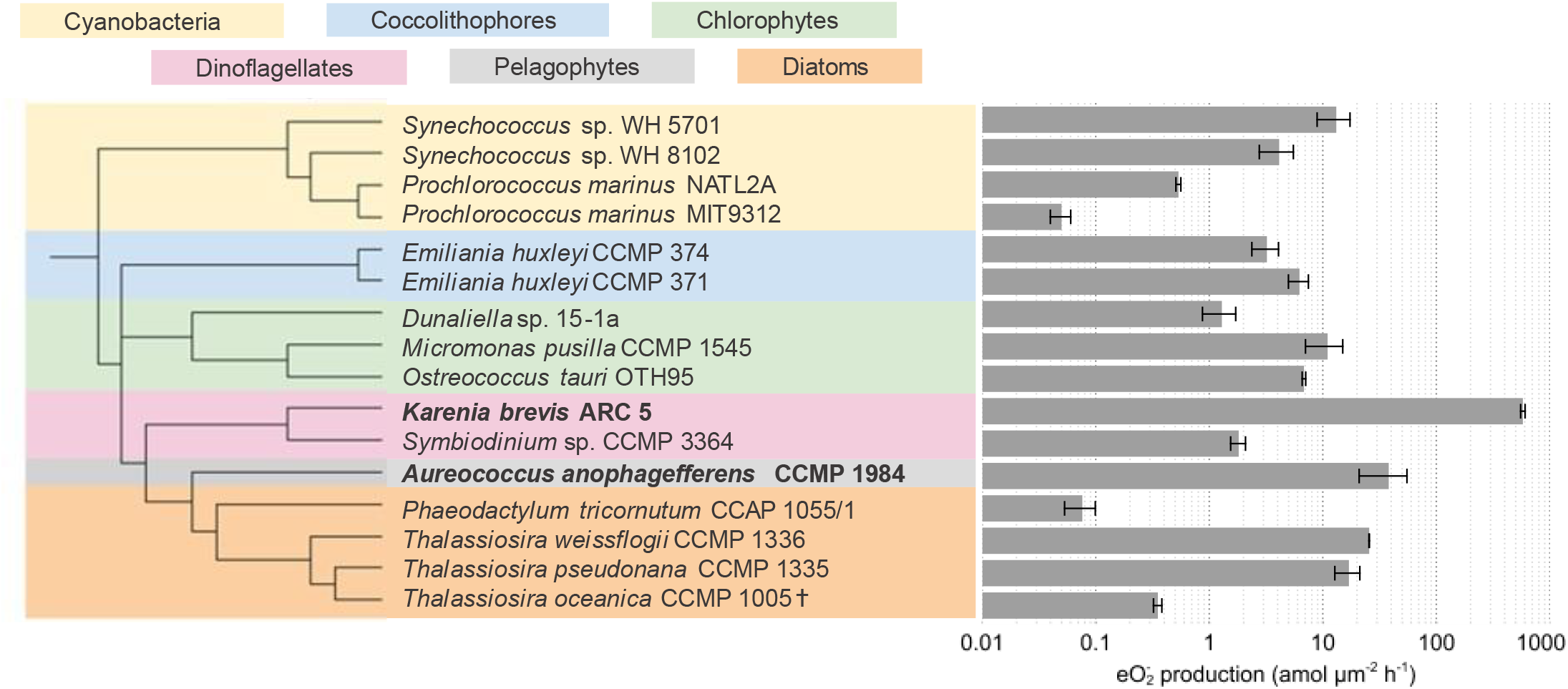
Surface area-normalized eO_2_^−^ production. Measurements were conducted under typical growth irradiance for each strain (see Supplementary Table 1). Leaf colors on phylogenetic tree indicate the different taxonomic groups. Bold font indicates harmful algal bloom-forming strains. Cell surface areas are shown in Supplementary Table 5. † indicates data from [14].

In a set of cultivation experiments, cultures were grown with the addition of the flavoenzyme inhibitor diphenyl iodonium (DPI), which irreversibly binds to flavin adenine dinucleotide (FAD)/ flavin mononucleotide (FMN) and consequently blocks their electron transfer activity [19]. Treatments included DPI (2 μM and 0.02 μM final concentrations) dissolved in 10% DMSO, 10% DMSO (0.03% v/v final concentration), or no treatment (unamended). The DPI and DMSO treatments (75 μL each) were added once after exponential growth phase had begun. In these experiments, growth was monitored by obtaining daily fluorescence for *P. marinus* strains or by obtaining daily cell abundance for all other strains as detailed above. Specific growth rate (d^−1^) was calculated by finding the slope of the regression of the natural log-normalized cell abundance or fluorescence over time during exponential phase. Exponential phase was defined as the natural log-linear (R^2^ ≥ 0.98) portion of cell abundance or fluorescence over time.

### Production of eO_2_^−^

Production of eO_2_^−^ by phytoplankton cells was measured using the flow-through FeLume (II) analytical system (Waterville Analytical, Waterville, Maine, USA) via reaction with the O_2_^−^-specific chemiluminescent probe methyl *Cypridina* luciferin analog (MCLA), as previously described [20–22]. To do so, cells were gently deposited onto an inline filter (0.22 μm for most strains, except 0.1 μm for cyanobacteria and smaller picoeukaryotes, polyethersulfone membrane, 13 mm diameter) using a syringe and continuously washed (2 mL min^−1^) with phosphate-buffered artificial seawater (20 mM phosphate; pH = 7.6) that matched the salinity of the culture media. The eO_2_^−^ within this cell-free effluent reacted with the MCLA reagent [4 μM MCLA, 0.1 M MES, 75 μM diethylenetriamine pentaacetic acid (DTPA); pH = 6] at the center of a spiral flow cell sitting below a photomultiplier tube housed within a light-tight box. Chemiluminescent data emitted from the reaction between eO_2_^−^ and MCLA were collected in real time using Waterville Analytical software. Autooxidation of the MCLA reagent generates O_2_^−^. Therefore, chemiluminescence was measured from cell-free blanks (i.e., MCLA reagent + artificial seawater) just prior to depositing phytoplankton cells onto the inline syringe filter for each analysis. These blank chemiluminescent signals were subtracted from the subsequent biological chemiluminescent signals (i.e., MCLA reagent + artificial seawater + cells). A steady-state signal was obtained by allowing chemiluminescent signals of both blanks and cells to stabilize (≤ 5% CV) for at least 1 min. Superoxide dismutase (SOD; final concentration of ~800 U L^−1^) was added at the end of each analysis to confirm chemiluminescent signals were due to O_2_^−^.

FeLume calibrations were performed with standard additions of potassium superoxide (KO_2_), as previously described [21, 22]. First, primary KO_2_ stocks were prepared by dissolving in a basic solution (0.03 N NaOH, pH=12.5; 90 μM DTPA). Then, absorbance of primary KO_2_ stocks [23] was measured at 240 nm before immediately being added to artificial seawater. The decay of O_2_^−^ was measured in this solution on the FeLume. The primary KO_2_ stocks were again measured at an absorbance of 240 nm after addition of SOD (~ 800 U L^−1^, final concentration). Absorbance measurements of KO_2_ primary stocks before and after SOD addition were used to quantify the O_2_^−^ concentrations by applying the extinction coefficient of O_2_^−^ corrected for production of H_2_O_2_ at 240 nm and a pH of 12.5 (2183 L mol^−1^ cm^−1^). Biological chemiluminescent signals were converted to steady-state eO_2_^−^ concentrations by dividing the chemiluminescent signals by the sensitivity of the analysis (chemiluminescent counts pM^−1^) obtained through standard addition calibrations. Then eO_2_^−^ concentrations (pM) were converted to production rates by multiplying by the flow rate (1 or 2 mL min^−1^), dividing that number by the number of cells loaded on the inline syringe filter, and converting final units to amol cell^−1^ h^−1^ or amol μm^−2^ h^−1^ to normalize to cell surface area. To ensure the same number of healthy cells was analyzed between replicates during an experiment, a cell concentration was first obtained via flow cytometry as detailed above. These production rates account for simultaneous production and decay of eO_2_^−^ and are therefore net production rates.

Production of eO_2_^−^ by phytoplankton cells was measured in the presence of DPI or the solvent control, DMSO. To do so, the artificial seawater (20 mM phosphate; pH = 7.6) that continuously washes cells during analysis was amended daily by heating and stirring for one hour, adding 20 μM DPI dissolved in 0.3% DMSO, or adding 0.3% DMSO alone (final concentrations), and allowing the solution to cool to room temperature. For these experiments, eO_2_^−^ measurements were collected at the growth irradiance of each phytoplankton stain (Supplementary Table 1). Irradiance was emitted from a dimmable soft white LED light bulb controlled with a manual dimmer and placed directly above the sample. A standard light bulb, which emits light in the visible range and little ultraviolet irradiation compared to natural sunlight [24], rather than a full spectrum light source was used to minimize abiotic ROS production while stimulating photosynthesis [9]. Irradiance was monitored with a micro quantum photosynthetically active radiation (PAR) sensor (Walz, Effeltrich, Germany). Data analysis and calculation of eO_2_^−^ production rates in the presence of DPI followed that of Plummer et al. [22] and is detailed above.

In another set of experiments, eO_2_^−^ production was measured as a function of increasing irradiance from ambient light levels (3-6 μmol m^−2^ s^−1^) to ~2250 μmol m^−2^ s^−1^ using a dimmable soft white LED light bulb, as previously described [14]. Then, these eO_2_^−^ production rates 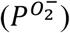 and irradiance (*E*) data were fit to an equation modified for eO_2_^−^ production from the double exponential photosynthesis-irradiance (PI) model by Platt et al.[25] following the methods of Diaz et al. [14]:

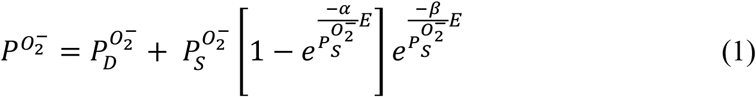

Where the fitted constants included α, the initial linear slope of the best fit curve; *β*, the parameter describing the decrease of eO_2_^−^ at high irradiances; 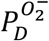,the net production rate of eO_2_ ^−^ in the dark; and 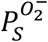,an estimate of the light-saturated rate of net eO_2_^−^ production if β=0.

Light-saturated rates of net eO_2_^−^production 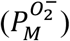 and the minimum saturating irradiance of net eO_2_^−^ production 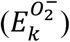were calculated by the following equations:

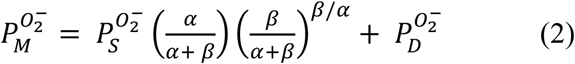

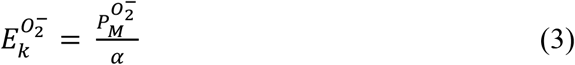

Taxon-specific constants are reported in Supplementary Table 2.

### Photophysiology

Photophysiology was monitored using either a Satlantic fluorescence induction and relaxation (FIRe) fluorometer system (Sea-bird Scientific, Bellevue, Washington, USA) or a pulse amplitude modulation (WATER-PAM) fluorometer (Walz, Effeltrich, Germany) similar to Diaz et al. [14]. Prior to measurements, phytoplankton were incubated with 2 μM DPI dissolved in 0.2% DMSO or 0.2% DMSO (final concentrations) for at least 30 min either at their growth irradiance (Supplementary Table 1) or in dark conditions. Photochemical efficiency of photosystem II (PSII) was determined by calculating photosynthetic efficiency (F_v_/F_m_) in the dark-adapted or light adapted state using the equation:

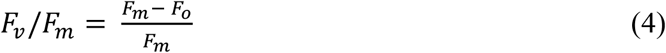

where *F*_*m*_ is the maximum fluorescence yield, and *F*_*o*_ is the minimum fluorescence yield. Decreases in F_v_/F_m_ correspond to increases in photoinhibition [26, 27].

### Rates of eO_2_^−^ production in the North Pacific Subtropical Gyre

We used our culture results to determine dark and light-driven rates of biological eO_2_^−^ production in the North Pacific Subtropical Gyre (NPSG). First, we calculated *in situ* irradiance levels (*E*_*z*_; μmol m^−2^ s^−1^) using an equation that describes how light intensity decreases exponentially with depth:

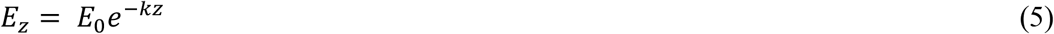

Where *E*_0_is defined as the irradiance level at the ocean surface obtained from seasonal PAR ranges in the NPSG [28] (324 to 613 μmol m^−2^ s^−1^ with a median of 481 μmol m^−2^ s^−1^), *k* is the average light attenuation coefficient throughout the seasons (0.04 m^−1^) [28], and *z* is depth (m). We used the long-term Hawaii Ocean Time-series (HOT) station ALOHA as a representative site in the NPSG. Depth (*z*) was defined as the mixed layer depth (MLD) unless otherwise noted. MLD was determined from the publicly available HOT dataset [29] over all years and seasons at station ALOHA. We defined the 5^th^ and 95^th^ percentiles (27 m – 109 m) as the range of typical MLD values, and we determined the median MLD to be 55 m (Supplementary Table 3). We assumed a MLD shallowing of 20 m in this region by the year 2200 based on the study by Luo and Rothstein [30], and therefore defined the median future MLD to be 35 m (Supplementary Table 4).

*In situ* biological eO_2_^−^ production rates (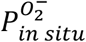; nmol L^−1^ d^−1^) can be described as a function of *in situ* irradiance levels (*E*_*z*_, determined by equation 5) using a modified form of equation (1):

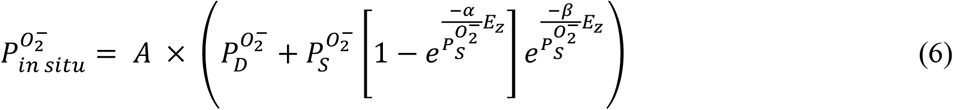

where, 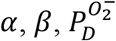,and 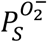 are taxon-specific constants determined from equation 1 (Supplementary Table 2) and *A* is the taxon-specific cell abundance in the environment (cells L^−1^; Supplementary Table 3). For these calculations, we utilized data from oligotrophic (e.g., *Synechococcus* sp. WH8102 instead of WH5701; *T*.*oceanica* instead of *T. pseudonana*) and environmentally relevant representatives (calcifying *E. huxleyi* CCMP 371 instead of noncalcifying CCMP 374). We assumed that cell abundances in the future would not change. Rates of dark biological eO_2_^−^ production were calculated using *E*_*z*_ = 0.

### Statistical analyses

All statistical analyses were performed in JMP Pro statistical software (SAS Institute Inc.,) or Microsoft Excel. A paired Student’s t-test (two-sample, two-tailed) and an unpaired, Student’s t-test (two-sample, two-tailed) were used to determine the effect of DPI on eO_2_^−^ production rates and phytoplankton growth rates, respectively. A Tukey-Honest Significance Difference (HSD) test was used to determine the effect of DPI on efficiency of PSII in light adapted and dark-adapted states. The significance threshold (alpha) was set to 0.05 for all statistical analyses.

## Results and Discussion

### Superoxide production varies widely among strains

We measured eO_2_^−^ production rates from 16 strains of prokaryotic and eukaryotic phytoplankton spanning a diversity of taxonomic groups and ecotypes (Fig. 1; Supplementary Table 1). These phytoplankton included representatives of the numerically dominant cyanobacteria *Synechococcus* and *Prochlorococcus*; *Emiliania huxleyi* – a widespread coccolithophore found from the equator to subpolar regions [31]; ecologically diverse green algae from the chlorophyte group; three species of *Thalassiosira*, one of the most abundant genera of diatoms [32]; and harmful bloom-forming algae from the dinoflagellate and pelagophyte groups. Since cell size influences eO_2_^−^ production rates [20, 33, 34], we normalized production rates to cell surface area (Supplementary Table 5). Production of eO_2_^−^ varied widely among the strains surveyed (Fig. 1 and Supplementary Fig. 2). For example, production rates of eO_2_^−^ measured under typical growth irradiances for each strain (see Supplementary Table 1) spanned 10^−3^ – 10^2^ amol μm^−2^ h^−1^; however, most phytoplankton strains produced eO_2_^−^ in the 10^−1^ – 10^1^ amol μm^−2^ h^−1^ range (Fig. 1 and Supplementary Fig. 2). Similar to previous results showing substantial eO_2_^−^ production by harmful algal species [35], the two highest eO_2_^−^ producers were the harmful bloom-forming algae *K. brevis* ARC 5 and *Aureococcus anophagefferens* CCMP 1984. While *A. anophagefferens* produced similar amounts of eO_2_^−^ to several non-harmful strains (10^1^ amol μm^−2^ h^−1^), *K. brevis* produced an order of magnitude more. *Prochlorococcus* (strain MIT9312) had the lowest rate of eO_2_^−^ production, consistent with previous observations [36]. Intraspecific variation was observed in *Prochlorococcus*, where *P. marinus* NATL2A produced 10-fold more eO_2_^−^ than *P. marinus* MIT9312. This finding may be related to the broader genomic capabilities of NATL2A, which contains more high-light induced photoprotective proteins than MIT9312 [37, 38].

**Fig. 2.**
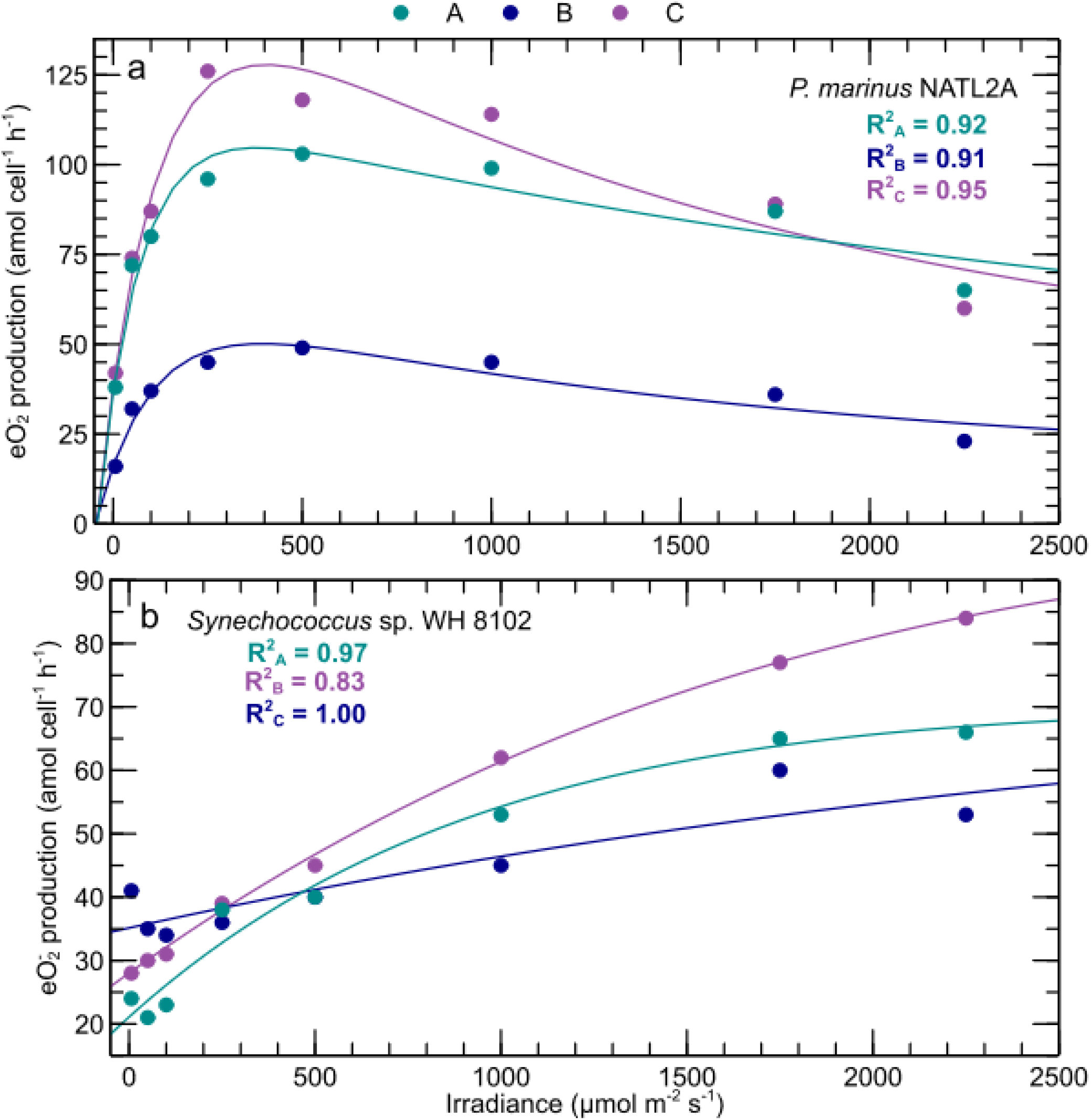
Irradiance versus eO_2_^−^ production. *P. marinus* NATL2A (a) shows the typical photoinhibition response observed in 15 of 16 strains studied, and *Synechococcus* sp. WH 8102 (b) shows an atypical response lacking photoinhibition. Irradiance and eO_2_^−^ production rate data were fit with a modified Photosynthesis-Irradiance model [14] (lines). See Materials and Methods and Supplementary Table 2. Each color represents a different biological replicate. Additional strains are shown in Supplementary Fig. 3.

### Irradiance drives superoxide production

Production of eO_2_^−^ has been attributed to photosynthesis in previous studies, usually based on eO_2_^−^ production measurements conducted at a limited number or range of light levels [39–42]. To expand on these observations, we measured and modeled eO_2_^−^ production across a wide range of irradiances. In all phytoplankton tested, irradiance drove eO_2_^−^ production (Fig. 2 and Supplementary Fig. 3). Typically, eO_2_^−^ production followed a pattern similar to a photosynthesis-irradiance model [14] (Supplementary Table 2; avg. ± SD of R^2^ in all strains = 0.95 ± 0.05). Consistent with the expected light-driven regulation of photosynthetic rates, eO_2_^−^ production rates increased linearly at low irradiances, became saturated at intermediate irradiances, and decreased at highest irradiances via photoinhibition (Supplementary Fig. 3 and exemplified by *P. marinus* NATL2A in Fig. 2a). The single exception to this trend was observed in the oligotrophic *Synechococcus* strain (WH8102), in which eO_2_^−^ production was not photoinhibited, yet still increased as a function of light (Fig. 2b). These findings support the hypothesis that photosynthetic processes regulate eO_2_^−^ in diverse phytoplankton species.

**Fig. 3.**
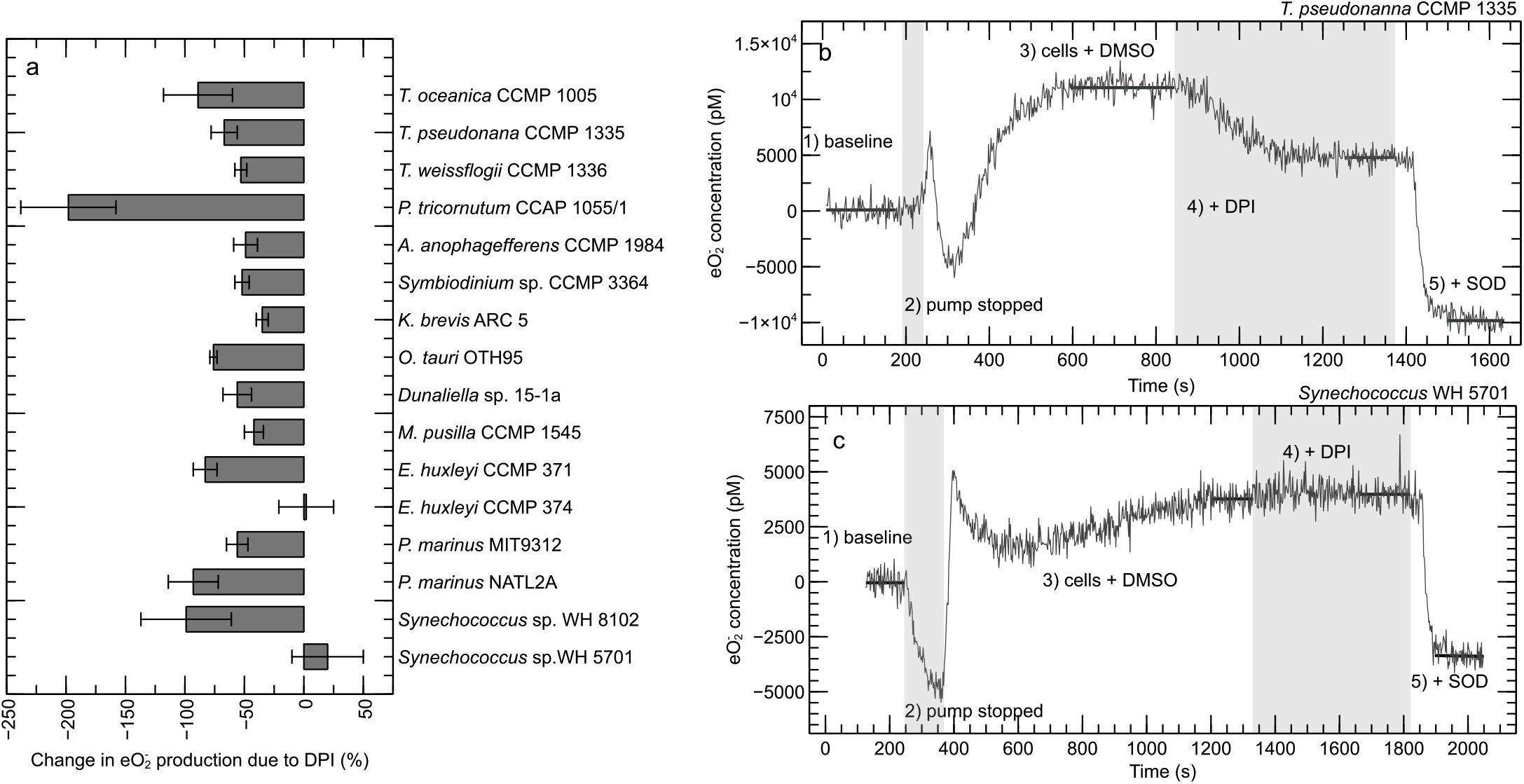
eO_2_^−^ production in the presence or absence of the flavoenzyme inhibitor DPI. Percent change in eO_2_^−^ production due to application of DPI (a). Error bars show standard deviation of the mean (n=3 biological replicates). (b,c) Time series of biological eO_2_^−^ production showing the typical inhibiting effect of DPI observed in 14 of 16 strains and exemplified by *T. pseudonanna* CCMP 1335 (b), and the atypical result showing no effect of DPI exemplified by *Synechococcus* sp. WH 5701 (c). Traces are split into five regions of analysis, as indicated (see Materials and Methods). Horizontal bars show stable signals from which the production rates were calculated. DPI was dissolved in DMSO, which had no impact on eO_2_^−^ production, as expected [14]. Negative eO_2_^−^ concentrations in the presence of superoxide dismutase (SOD) and changes in eO_2_^−^ production that are < −100% reflect degradation of background eO_2_^−^ originating from the analytical reagent (see Materials and Methods). Data for *T. oceanica* CCMP 1005 are from Diaz et al. [14]

### Flavoenzymes mediate eO_2_^−^ production

Superoxide (pKa=4.8) is an anion at typical cellular pH and has a short intracellular lifetime (~10^−5^ s) [43]. Therefore, eO_2_^−^ does not readily cross the plasma membrane that separates the cell from its surrounding environment and is unlikely to arise from passive diffusion or cell lysis [44]. Instead, biological eO_2_^−^ production is typically mediated by cell surface enzymes that catalyze electron transport across the plasma membrane [43, 44]. In phytoplankton, this transplasma membrane electron transport is thought to be driven by flavoenzymes that oxidize the cellular reductant NADPH, including NADPH oxidase (NOX) [14, 35, 41, 45–47] and glutathione reductase [14, 48], which are inhibited by diphenyl iodonium (DPI) [19, 48].

Here, we applied DPI and found that it inhibited eO_2_^−^ production in 14 of the 16 phytoplankton strains examined (Fig. 3a and Supplementary Fig. 2), consistent with a predominant role for flavoenzymes. Indeed, DPI quenched eO_2_^−^ production on a timescale of minutes in many strains (exemplified by *T. pseudonanna* CCMP 1335 in Fig. 3b). Yet here and in other studies [14, 41, 45-47], DPI did not completely eliminate eO_2_^−^ production in most organisms (Fig. 3a and Supplementary Fig. 2), suggesting that enzymes in addition to the flavoenzyme group are involved in eO_2_^−^ production. In the coccolithophore *E. huxley*i CCMP374 and the coastal *Synechococcus* strain WH5701, eO_2_^−^ production was insensitive to DPI, indicating that flavoenzymes may not produce eO_2_^−^ in these organisms (Fig. 3 and Supplementary Fig. 2).

### Flavoenzyme/eO_2_^−^ inhibitor impairs photosynthetic health and viability

Given the confirmed photoprotective role of eO_2_^−^ production in *T. oceanica* [49], we hypothesized that DPI would impair photosynthetic health in most phytoplankton that exhibited DPI-inhibitable eO_2_^−^ production. Therefore, we examined the effect of DPI on the efficiency of photosystem II (F_v_/F_m_) under light and dark conditions on a timescale of tens of minutes (see Materials and Methods). F_v_/F_m_ represents the fraction of total absorbed light energy that is used in photosynthesis, with higher levels typically representing better photosynthetic health. We found that DPI inhibited F_v_/F_m_ in 14 of 16 strains and that this inhibition was dependent on the presence of light (Fig. 4. groups Ia-c). In the majority of strains, DPI decreased F_v_/F_m_ under light-adapted conditions but not dark-adapted conditions (group Ia), while in other strains, DPI decreased F_v_/F_m_ under both light and dark conditions (group Ib). In two strains, DPI inhibited F_v_/F_m_ under light conditions yet stimulated F_v_/F_m_ under dark conditions (group Ic). Overall, DPI inhibited eO_2_^−^ production and F_v_/F_m_ in the majority (13/16) of strains (Fig. 4), consistent with the hypothesized role for eO_2_^−^ production in supporting phytoplankton photophysiology.

**Fig. 4.**
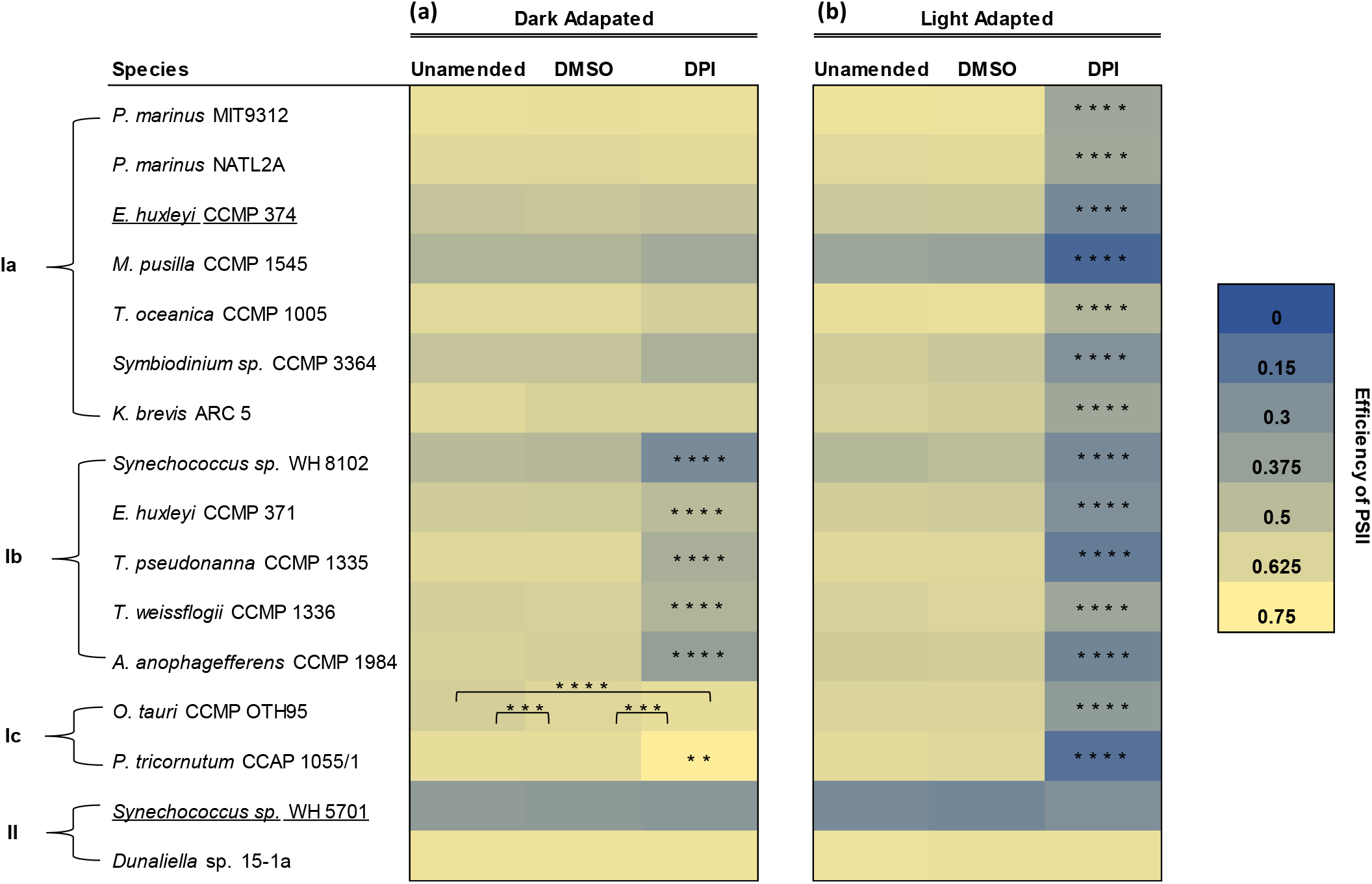
Efficiency of PSII in the presence or absence of the flavoenzyme inhibitor DPI. Phytoplankton were acclimated to (a) their growth irradiance (light adapted) or (b) darkness (dark adapted). All measurements were conducted in 0.2% DMSO, except the unamended control. Significant differences of the mean F_v_/F_m_ (n = 3 biological replicates) between treatments were found using a Tukey-Honest Significance Difference (HSD) test. P-values are indicated by asterisks, where **, ***, and **** signifies a p-value of <0.01, <0.001, and <0.0001, respectively. Responses of efficiency of PSII to DPI are categorized into four groups: Ia, inhibition under light adaptation only; Ib, inhibition under both light and dark adaptation; Ic, inhibition under light adaptation, yet stimulation under dark conditions; and II, no effect. Strains that are underlined did not show DPI-inhibitable eO_2_^−^ production. Data for *T. oceanica* CCMP 1005 are from Diaz et al. [14]

Based on the negative effect of DPI on eO_2_^−^ production and photosynthetic health in the majority of phytoplankton tested, we hypothesized that DPI would also impair viability. To test this hypothesis, we added two concentrations of DPI to early exponential phase cultures and compared their growth to DMSO and unamended controls (Supplementary Fig. 4). DPI inhibited growth and ultimately led to cell death in 15 out of 16 strains (Fig. 5 and Supplementary Fig. 5). Further, the effect of DPI was concentration-dependent in the majority of strains (group Ia), though some were only sensitive to the higher concentration of DPI (group Ib). Overall, DPI inhibited both eO_2_^−^ production and growth in the majority (14/16) of strains (Fig. 5), consistent with the hypothesized role for eO_2_^−^ production in promoting phytoplankton viability.

**Fig. 5.**
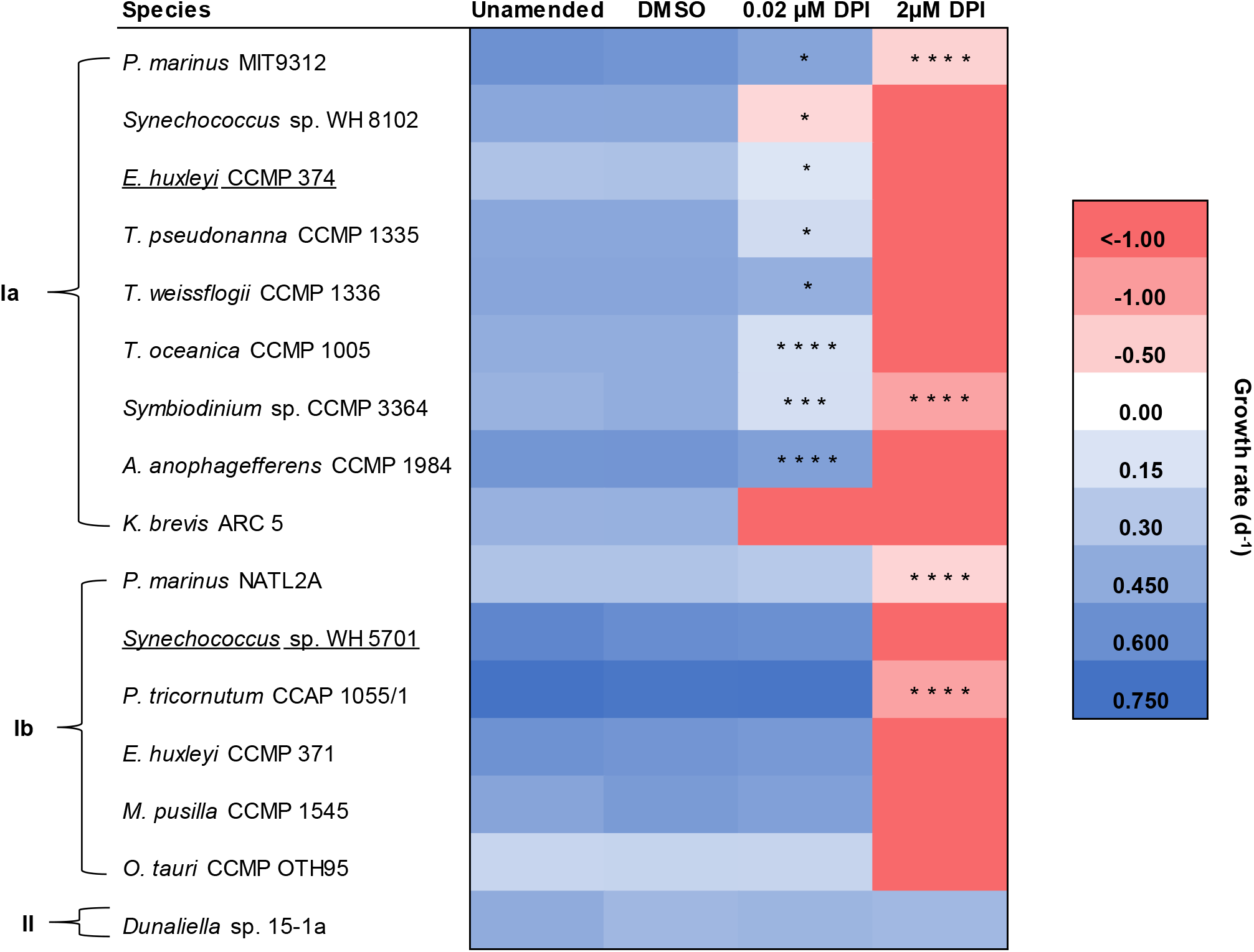
Specific growth rates in the presence or absence of the flavoenzyme inhibitor DPI. All cultures were grown in 0.03% DMSO, except the unamended control. Negative growth rates indicate culture death. Growth rates < −1.00 d^−1^ indicate that cells died at a rate that was faster than our limit of detection. Asterisks indicate significant differences of mean growth rates > −1.00 (n = 3 biological replicates) between the DPI and DMSO treatments using a Student’s t-test (unpaired, two-sample). Statistical results between all other conditions are shown in Supplementary Fig. 5. P-values are indicated by asterisks, where *, **, ***, and **** signifies a p-value of <0.05, <0.01, <0.001, and <0.0001, respectively. Responses of growth rates to DPI are categorized into 3 groups: Ia, significant declines in both the 0.02 μM DPI and 2μM DPI treatments; Ib, significant declines in only the 2 μM DPI treatment; and II, no effect. Strains that are underlined did not show DPI-inhibitable eO_2_^−^ production.

### Extracellular superoxide production supports phytoplankton health

In summary, for the 14 organisms showing DPI-inhibitable eO_2_^−^ production, DPI treatment also decreased photosynthetic efficiency and growth, eventually leading to cell death in all but one case (Table 1). Combined with the tight regulation of eO_2_^−^ production by photosynthesis, these findings are consistent with the hypothesis that eO_2_^−^ production supports the health of most phytoplankton by protecting against photodamage. It is unlikely that our findings could be explained by a broad and non-specific cytotoxicity of DPI because adverse health outcomes were not always linked to the inhibition of eO_2_^−^ production by DPI and vice versa, such as in *Dunaliella* sp. 15-1a, *Synechococcus* sp. WH5701, and *E. huxleyi* CCMP374 (Table 1). Also, our photophysiological experiments show that for most strains, DPI was only toxic to photosynthetic health in the light and not in the dark (e.g., Fig. 4 groups 1a and 1c), which is consistent with the proposed light-dependent eO_2_^−^ photoprotection strategy.

Exceptions to the general trend we observed included the extremophile *Dunaliella* sp. 15-1a, in which DPI inhibited eO_2_^−^ production to a similar degree as other phytoplankton (Fig. 3), but DPI did not affect photosynthetic health or growth (Figs. 4, 5 and Table 1), demonstrating that eO_2_^−^ production may not serve a photoprotective mechanism in this species. We could not test the potential photoprotective role of eO_2_^−^ production in *Synechococcus* sp. WH 5701 or *E. huxleyi* CCMP 374 because eO_2_^−^ production by these organisms was insensitive to DPI (Fig. 3). However, we did observe some impairments to health from DPI treatments with these strains (Figs. 4, 5), consistent with the diverse physiological roles of flavoenzymes [50]. For instance, in *E. huxleyi* CCMP 374, DPI inhibited growth and photosynthetic health, whereas DPI only inhibited growth in *Synechococcus* sp. WH 5701 (Table 1).

### Superoxide production aids redox balance

Phytoplankton commonly employ energy-dissipating strategies to avoid over-reduction and damage of the photosynthetic apparatus, such as nonphotochemical quenching [51], cyclic electron flow [52], and the Mehler reaction [53]. The process of making eO_2_^−^ may serve a similar photoprotective role. During photosynthesis, ATP and the reductant NADPH are produced. Both ATP and NADPH are then consumed in biosynthetic pathways, such as the Calvin cycle for carbon fixation. The ATP:NADPH ratio of photosynthesis (~1.3) is estimated to be lower than what is needed to fuel the Calvin cycle (1.5) [54, 55], suggesting a relative “excess” of NADPH. Moreover, the ratio of NADPH:NADP^+^ increases with increasing light levels [56], which could contribute to redox imbalance if NADPH is not used or recycled to NADP^+^. Because NADPH is the substrate used by enzymes that generate eO_2_^−^ [57] producing eO_2_^−^ is one way to recycle excess cellular NADPH to NADP^+^, thereby preventing overreduction. Similar to other photoprotective pathways, our results are consistent with eO_2_^−^ production helping prevent reductive stress by regulating cellular NADPH levels, promoting intracellular redox balance, and supporting health in many diverse phytoplankton.

Outside of photosynthesis, eO_2_^−^ production may play an even broader role in biological redox homeostasis. The majority of biotic eO_2_^−^ production rates in laboratory and field settings have been measured in the absence of light to eliminate abiotic O_2_^−^ production via photochemical reactions [20, 36, 58, 59]. Indeed, phytoplankton produce eO_2_^−^ under dark conditions, but at lower rates than in the light [14, 22, 41, 42] (Fig. 2 and Supplementary Fig. 3). In the dark, these microbes likely produce eO_2_^−^ using light-independent sources of reducing power, such as NADPH derived from the oxidative pentose phosphate pathway. Yuasa et al. reported that the harmful bloom-forming microalga *Chattonella antiqua* produces elevated levels of eO_2_^−^ to regulate imbalanced NADPH:NADP^+^ ratios influenced by the oxidative pentose phosphate pathway that occur under nitrogen and phosphorous limitation [60]. In these ways, eO_2_^−^ production appears to be a cellular redox acclimation strategy that is controlled by nutrient availability, as well as light exposure in phytoplankton.

### Implications for light-driven superoxide production in the marine environment

Environmental rates of biological superoxide production are not fully understood because most observations have been conducted only under dark conditions. Our results advance the current understanding of biological superoxide production in the marine environment by illuminating potential dynamics in the daytime. For example, we can extrapolate the eO_2_^−^ - irradiance response of each phytoplankton strain to the environment by accounting for its *in situ* cell abundance and level of light exposure, including darkness (see Materials and Methods). The sum of these taxon-specific rates represents eO_2_^−^ production by the phytoplankton community.

We applied the above approach to Station ALOHA, a long-term time series site in the North Pacific Subtropical Gyre (NPSG). First, we estimated the phytoplankton-driven eO_2_^−^ production under dark conditions and compared our result to the only available measurements of biological eO_2_^−^ production at Station ALOHA, which were conducted in the dark [58]. Because half of our cultures including *Prochlorococcus* were axenic during our experiments (Supplementary Table 1), our calculation underestimates eO_2_^−^ degradation by heterotrophic bacteria, which would be significant in the environment [1]. Yet, our approach is conservative in the sense that we do not account for the influence of nutrient stress, which is likely prevalent in the oligotrophic NPSG, and would probably increase eO_2_^−^ production rates [60]. Nevertheless, our extrapolation of 9 nM d^−1^ at 0 μmol m^−2^ s^−1^ by the phytoplankton community agrees well with measurements by Roe et al. [58], who reported dark rates of eO_2_^−^ production at Station ALOHA that averaged between ~2.6 to 6 nM d^−1^. Due to the numerical dominance of cyanobacteria in the phytoplankton community at Station ALOHA (Supplementary Table 3 and references therein), *Prochlorococcus* and *Synechococcus* were the dominant contributors to eO_2_^−^ production under dark conditions, producing 87% and 11% of eO_2_^−^, respectively.

Next, we used a similar approach to determine the effect of light exposure on phytoplankton-driven eO_2_^−^ production rates at Station ALOHA. To do so, we used ocean surface measurements of photosynthetically active radiation (PAR) [28] to calculate *in situ* PAR fluxes throughout the mixed layer [29]. *Prochlorococcus* and *Synechococcus* generated 82% – 99% and 1-17% of phytoplankton-derived eO_2_^−^ across the range of light levels found throughout the mixed layer, respectively (see Materials and Methods). Additionally, based on data from Malmstrom et al.[61], we determined that the high-light adapted *Prochlorococcus* ecotypes (represented by MIT9312) constitute ~99-100% of the *Prochlorococcus* community within the mixed layer PAR range. Therefore, our subsequent analyses focused on *Prochlorococcus* MIT9312 based on its dominance. At the median annual PAR level and MLD, we estimate light-driven eO_2_^−^ production by *Prochlorococcus* to be 14 ± 7 nM d^−1^ (Supplementary Table 4), representing a ~50% increase over the modeled community eO_2_^−^ production rate under dark conditions. Our light-driven estimates of biological eO_2_^−^ production do not account for UV irradiance (see Materials and Methods), which has the potential to both diminish or amplify environmental O_2_^−^ production through its inhibitory effect on photosynthesis [63, 64], or through the abiotic photooxidation of organic matter [1], respectively. Nevertheless, our results indicate that PAR exposure has the potential to increase environmental rates of biological eO_2_^−^ production.

Light-driven eO_2_^−^ production by phytoplankton may have implications for future oceans impacted by climate change. For example, MLDs in the NPSG are predicted to shallow up to 20 m by the year 2200 [30]. We estimate that a 20 m shallowing of the historic median annual MLD at Station ALOHA would increase light-dependent eO_2_^−^ production to 21 ±10 nM d^−1^ using the median annual PAR level (see Materials and Methods; Supplementary Table 4). This change corresponds to a 50% increase in eO_2_^−^ production compared to the historic median annual MLD (Supplementary Fig. 6). This estimate is likely conservative, because we did not account for potential climate-driven changes in *Prochlorococcus* abundances, which are predicted to increase in this region [65]. Elsewhere in the global ocean, enhanced water column stratification has contributed to the deepening of ocean MLD worldwide [66]. And in polar regions, loss of sea ice has increased water column light exposure [67]. Phytoplankton photoacclimation is also predominant on a global scale [68]. Therefore, future work should consider how phytoplankton photoacclimation may alter environmental eO_2_^−^ levels, and the coupled cycling of oxygen, carbon, and metals, in response to climate-driven changes in water column structure and light exposure.

## Acknowledgements

This work was supported by a National Science Foundation Graduate Research Fellowship [2017250547] to S.P.; a University of California President’s Dissertation Year Fellowship to S.P.; a National Science Foundation Research Experiences for Undergraduates award [1659793]; a Sloan Research Fellowship from the Alfred P. Sloan Foundation to J.M.D. [FG-2019-12550]; and a Simons Early Career Investigator in Marine Microbial Ecology and Evolution Award from the Simons Foundation to J.M.D. [678537].

## Data Availability

The datasets generated during and/or analyzed during the current study are available from the corresponding authors on reasonable request.

## Author Contributions

J.M.D and S.P. conceived and designed the study. S.P. conducted the experiments with assistance from S.G., analyzed the data, and wrote the manuscript. All authors contributed to interpretation of the results and preparing the manuscript.

## Figure Legends

**Table 1. The effect of DPI on eO_2_^−^ production, growth, and photosynthetic efficiency (F_v_/F_m_) of phytoplankton studied**. Bold font indicates a HAB forming species. Effects of DPI include significant inhibition (-) or no effect (+/-); *results from Diaz et al. [14]

