## Supplementary Information for "Extracellular superoxide production is a widespread photoacclimation strategy in phytoplankton"

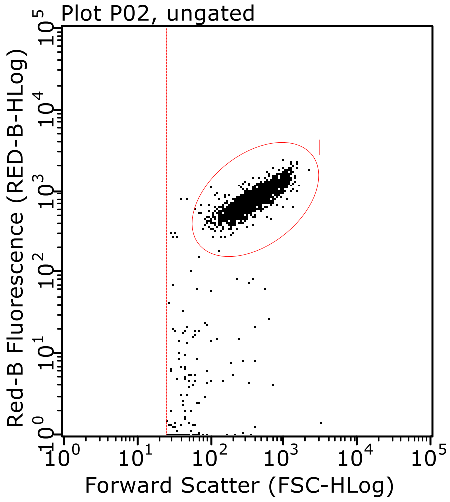


**Supplementary Fig. 1. Exemplary plot of flow cytometry gates used to enumerate phytoplankton**. Phytoplankton concentrations (cells $mL^{-1}$) were determined using diagnostic gates of red fluorescence versus forward scatter of exponentially growing monocultures. An exponentially growing population of *M. pusilla* CCMP 1545 is shown.

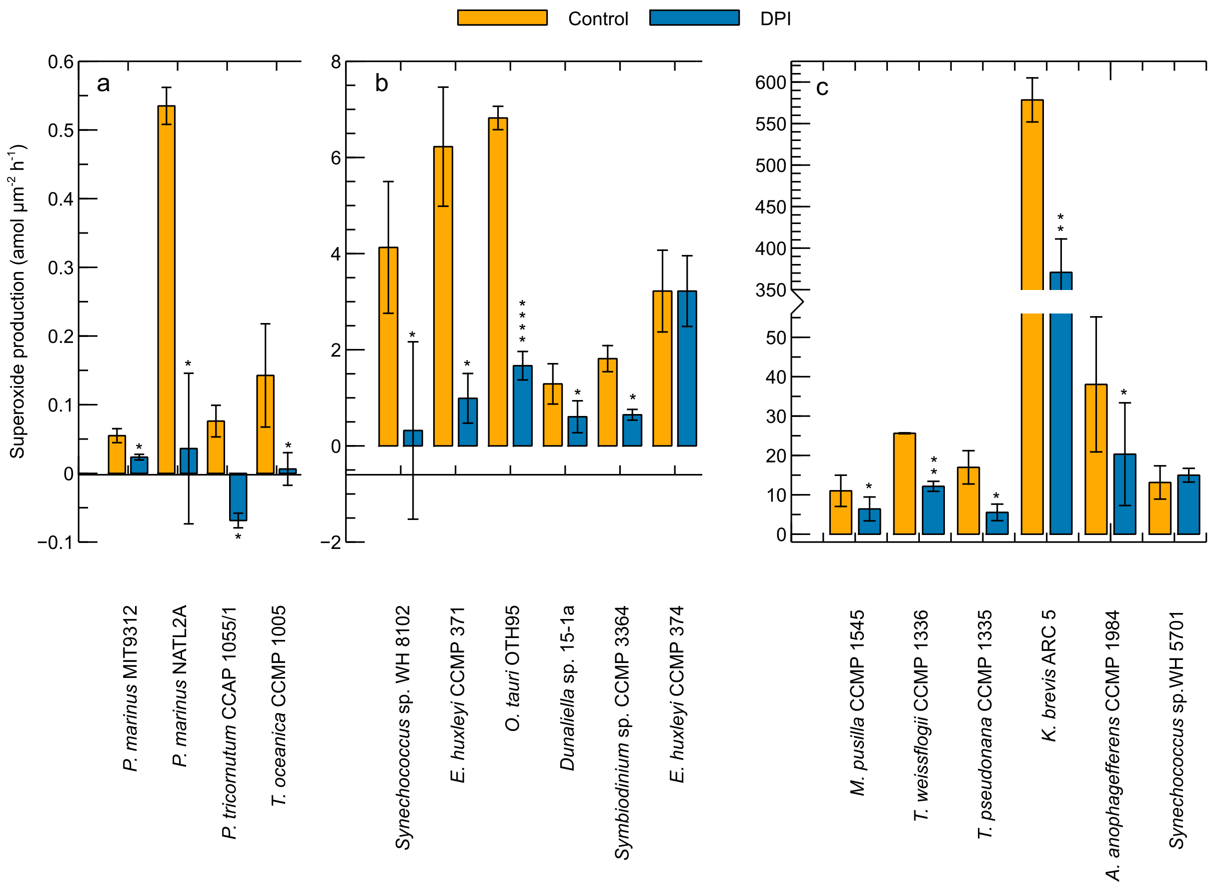


**Supplementary Fig. 2. Rates of $eO_2^-$ production in the presence and absence of the flavoenzyme inhibitor DPI.** All measurements were conducted in 0.3% DMSO. Production rates were normalized to cell surface area (Supplementary Table 5). Note the different y-axis scales on each panel. Significant differences of mean $eO_2^-$ production rates (n = 3 biological replicates) versus the DMSO control were found with a Student's t-test (paired, two sample). P-values are indicated by asterisks, where *, **, and **** signifies a p-value of <0.05, <0.01, and <0.0001, respectively. Error bars represent one standard deviation of the mean. Data from *T. oceanica* CCMP 1005 were taken from Diaz et al. [1].

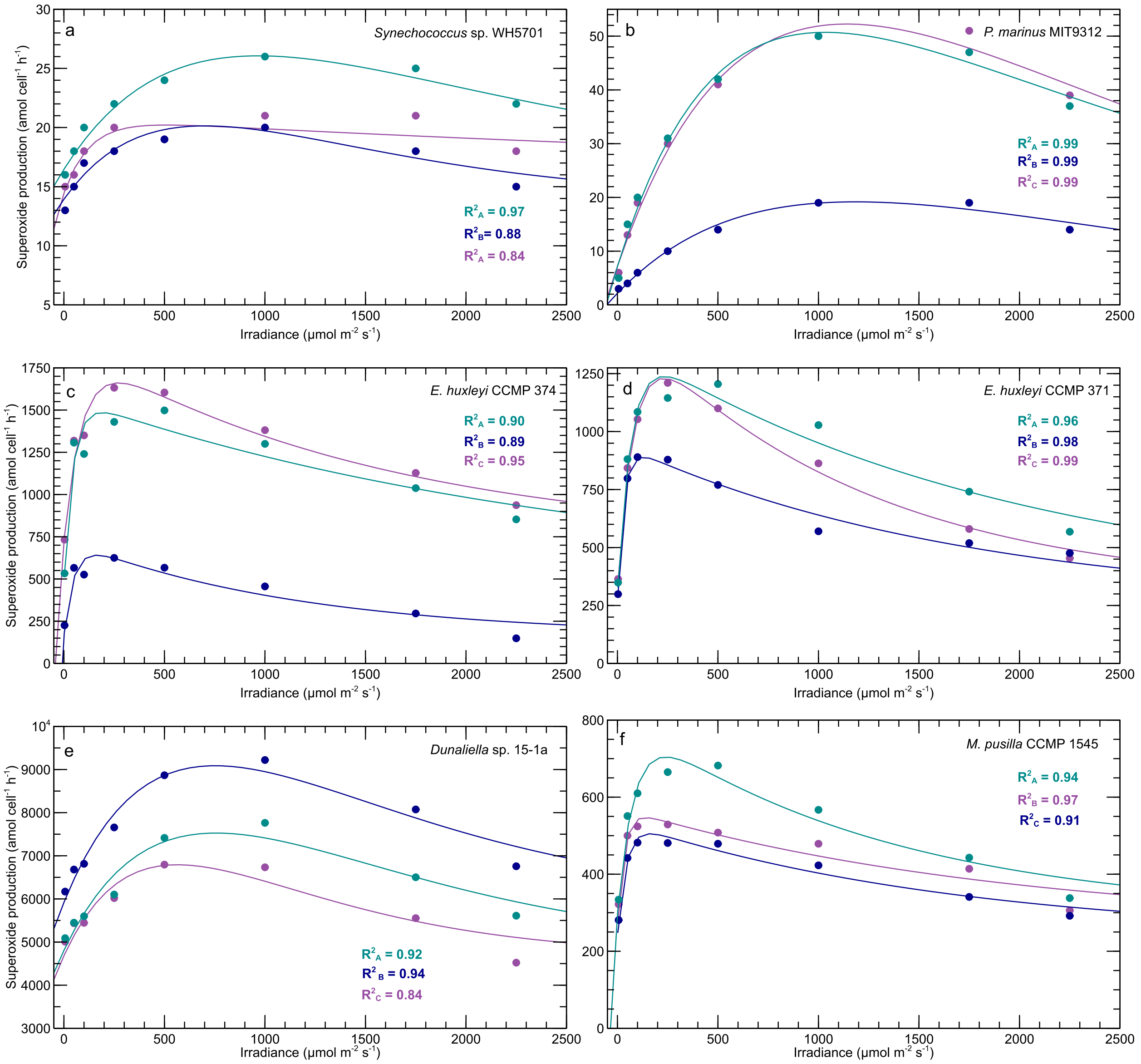

a
Synechococcus sp. WH5701
$R^2_A$ = 0.97
$R^2_B$= 0.88
$R^2_A$ = 0.84
b
P. marinus MIT9312
$R^2_A$ = 0.99
$R^2_B$ = 0.99
$R^2_C$ = 0.99
c
E. huxleyi CCMP 374
$R^2_A$ = 0.90
$R^2_B$ = 0.89
$R^2_C$ = 0.95
d
E. huxleyi CCMP 371
$R^2_A$ = 0.96
$R^2_B$ = 0.98
$R^2_C$ = 0.99
e
Dunaliella sp. 15-1a
$R^2_A$ = 0.92
$R^2_B$ = 0.94
$R^2_C$ = 0.84
f
M. pusilla CCMP 1545
$R^2_A$ = 0.94
$R^2_B$ = 0.97
$R^2_C$ = 0.91
Superoxide production (amol cell$^{-1}$ h$^{-1}$)
Irradiance (µmol m$^{-2}$ s$^{-1}$)

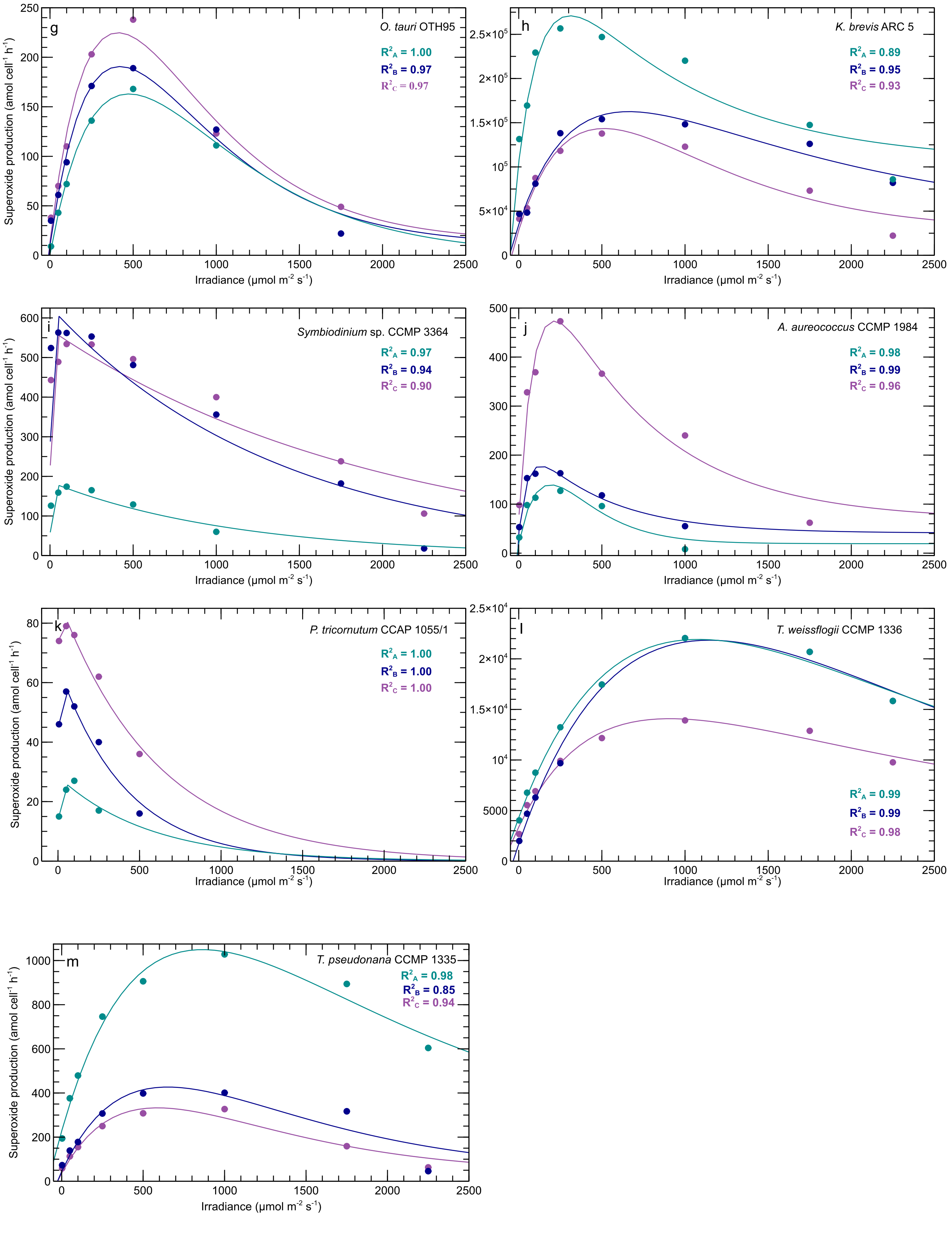
g
O. tauri OTH95
$R^2_A$ = 1.00
$R^2_B$ = 0.97
$R^2_C$ = 0.97
Superoxide production (amol cell$^{-1}$ h$^{-1}$)
Irradiance (µmol m$^{-2}$ s$^{-1}$)
h
K. brevis ARC 5
$R^2_A$ = 0.89
$R^2_B$ = 0.95
$R^2_C$ = 0.93
Irradiance (µmol m$^{-2}$ s$^{-1}$)
i
Symbiodinium sp. CCMP 3364
$R^2_A$ = 0.97
$R^2_B$ = 0.94
$R^2_C$ = 0.90
Superoxide production (amol cell$^{-1}$ h$^{-1}$)
Irradiance (µmol m$^{-2}$ s$^{-1}$)
j
A. aureococcus CCMP 1984
$R^2_A$ = 0.98
$R^2_B$ = 0.99
$R^2_C$ = 0.96
Irradiance (µmol m$^{-2}$ s$^{-1}$)
k
P. tricornutum CCAP 1055/1
$R^2_A$ = 1.00
$R^2_B$ = 1.00
$R^2_C$ = 1.00
Superoxide production (amol cell$^{-1}$ h$^{-1}$)
Irradiance (µmol m$^{-2}$ s$^{-1}$)
l
T. weissflogii CCMP 1336
$R^2_A$ = 0.99
$R^2_B$ = 0.99
$R^2_C$ = 0.98
Irradiance (µmol m$^{-2}$ s$^{-1}$)
m
T. pseudonana CCMP 1335
$R^2_A$ = 0.98
$R^2_B$ = 0.85
$R^2_C$ = 0.94
Superoxide production (amol cell$^{-1}$ h$^{-1}$)
Irradiance (µmol m$^{-2}$ s$^{-1}$)

**Supplementary Fig. 3. $eO_2^-$ production rates measured at increasing irradiances from triplicate batch cultures of model phytoplankton strains.** Irradiance and $eO_2^-$ production rate data (circles) were fit with a photosynthesis-irradiance model by Platt et al. [2] that was adapted by Diaz et al. [1] for $eO_2^-$ production rates (lines). Each color represents a different biological replicate. $R^2$ values of the model fit for each biological replicate are provided. Model rates are presented in Supplementary Table 2. Results for *T. oceanica* are from Diaz et al. [1] and are not presented here.

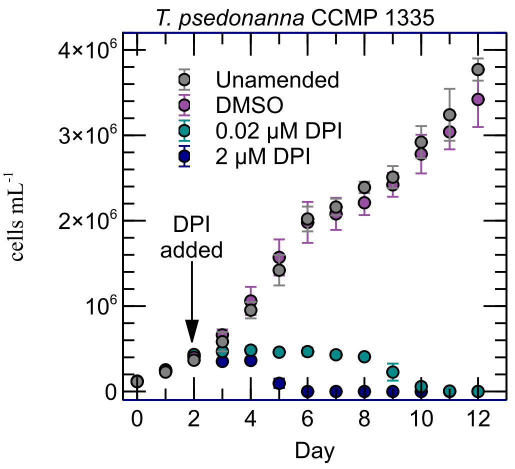


**Supplementary Fig. 4. The effect of DPI on growth of *T. pseudonana* CCMP 1335.** Error bars show standard deviation of the mean of biological replicates (n=3).

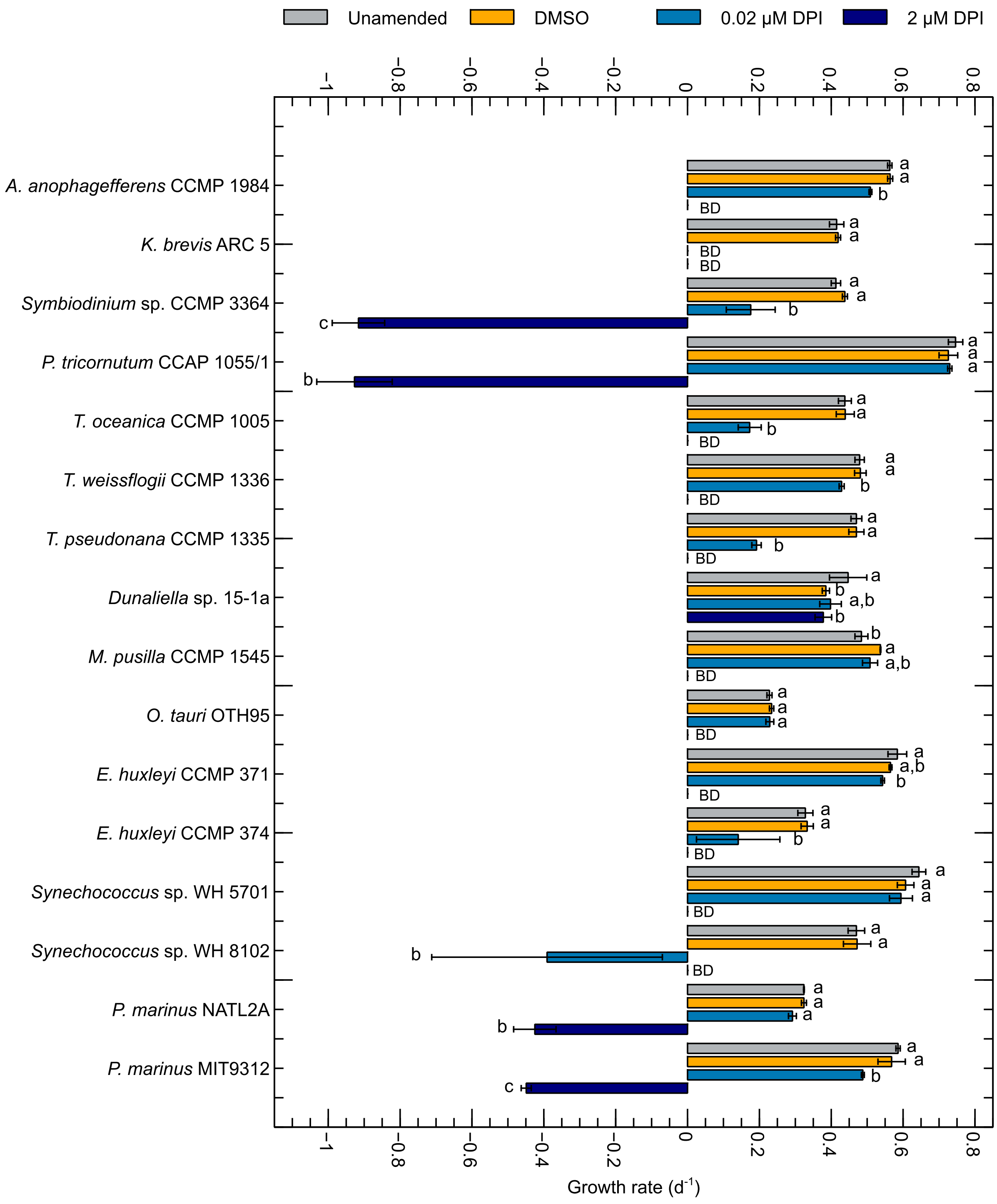
Unamended
DMSO
0.02 μM DPI
2 μM DPI
A. anophagefferens CCMP 1984
K. brevis ARC 5
Symbiodinium sp. CCMP 3364
P. tricornutum CCAP 1055/1
T. oceanica CCMP 1005
T. weissflogii CCMP 1336
T. pseudonana CCMP 1335
Dunaliella sp. 15-1a
M. pusilla CCMP 1545
O. tauri OTH95
E. huxleyi CCMP 371
E. huxleyi CCMP 374
Synechococcus sp. WH 5701
Synechococcus sp. WH 8102
P. marinus NATL2A
P. marinus MIT9312
Growth rate (d$^{-1}$)

**Supplementary Fig. 5. Specific growth rates in the presence or absence of the flavoenzyme inhibitor DPI.** All cultures were grown in 0.03% DMSO, except the unamended control. BD stands for below detection and indicates cell death at a rate that was faster than our limit of detection. Significant differences of mean growth rates > -1.00 ($n = 3$ biological replicates) between treatments were found using a Student's t-test (unpaired, two-sample). Treatments not connected by the same letter within a strain are significantly different ($p$-value $< 0.05$). Error bars show the standard deviation of the mean of biological replicates ($n=3$).

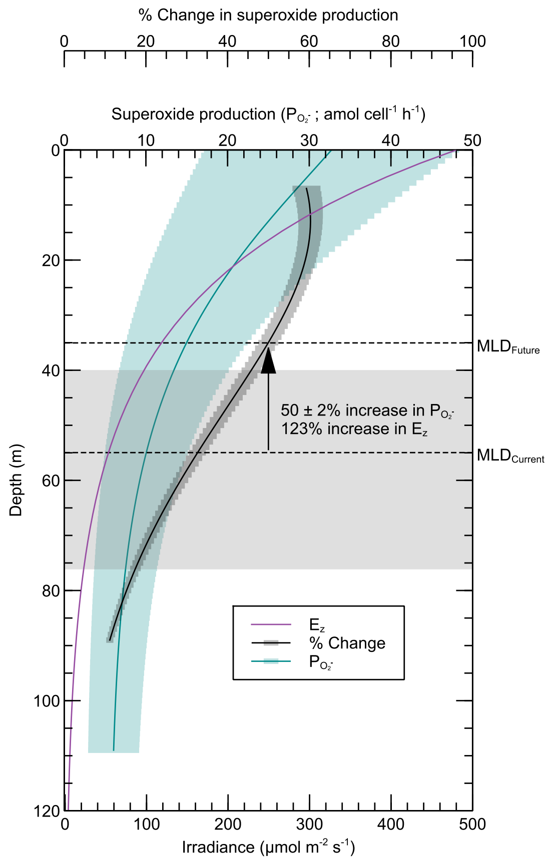


**Supplementary Fig. 6. Calculated production rates of $eO_2^-$ by *Prochlorococcus* ($P_{O2-}$) under current and future conditions in the North Pacific Subtropical Gyre.** Production rates of $eO_2^-$ by *Prochlorococcus* ($P_{O2-}$) and irradiance ($E_z$) as a function of depth in the North Pacific Subtropical Gyre (Station ALOHA) were based on median surface irradiance or $E_0$ [3] (see Materials and Methods). Dashed lines indicate median current ($MLD_{Current}$) and future ($MLD_{Future}$= $MLD_{Current}$ – 20 m) [4] mixed layer depths. Shaded area indicates the 25$^{th}$ and 75$^{th}$ percentile of the $MLD_{Current}$. Black line shows the % change in per-cell $eO_2^-$ production at each depth between the 5$^{th}$ and 95$^{th}$ percentile of the MLD under current and future conditions. Shaded bars around $P_{O2-}$ and % change show standard deviation of the mean of biological replicates (n=3). Using equation (5), we determined that a 20 m shoaling would increase $E_z$ at the MLD by 123% by the year 2200. For instance, a 20 m shoaling from 55 m to 35 m would increase $E_z$ by

~53 μmol $m^{-2}$ $s^{-1}$ to ~119 μmol $m^{-2}$ $s^{-1}$ or 123%. $E_z$ data do not have standard deviation because these data are based on a fundamental equation modeling the relationship between surface irradiance and depth (see Materials and Methods).

**Supplementary Table 1. Growth conditions, sources, and ecotypes of the 16 model phytoplankton strains selected for the study.** Strains are marine unless otherwise stated. L1 + Si [5], f/2 and f/2 + Si [6], SN [7], and Pro99 [8] media were prepared according to standard procedures.

| Class | Genus species | Strain | Media | Growth Irradiance (μmol $m^{-2}$ $s^{-1}$) | Temperature (°C) | Axenic | Ecotype Notes |
|---|---|---|---|---|---|---|---|
| Prymnesiophyceae | *Emiliania huxleyi* | CCMP 374 | f/2 | 100-130 | 18 | Yes | non-calcifying |
| | *Emiliania huxleyi* | CCMP 371 | f/2 | 100-130 | 18 | Yes | calcifying |
| Chlorophyceae | *Ostreococcus tauri* | OTH95 | f/2 | 100-130 | 18 | No | |
| | *Micromonas pusilla* | CCMP 1545 | f/2 | 100-130 | 18 | Yes | |
| | *Dunaliella* | 15-1a | L1 + Si (120 ppt) | 100-130 | 18 | No | hypersaline |
| Bacillariophyceae | *Thalassiosira pseudonana* | CCMP 1335 | f/2 + Si | 100-130 | 18 | No | coastal |
| | *Thalassiosira weissflogii* | CCMP 1336 | f/2 + Si | 100-130 | 18 | Yes | coastal |
| | *Thalassiosira oceanica* | CCMP 1005 | f/2 + Si | 100-130 | 23 | Yes | oligotrophic |
| | *Phaeodactylum tricornutum* | CCAP 1055/1 | L1 + Si | 100-130 | 18 | No | |
| Dinophyceae | *Symbiodinium* sp. | CCMP 3364 | L1 + Si | 100-130 | 23 | No | coral symbiont |
| | *Karenia brevis* | ARC 5 | L1 + Si | 100-130 | 23 | No | HAB-forming |
| Pelagophyceae | *Aureococcus anophagefferens* | CCMP 1984 | L1 + Si | 100-130 | 18 | No | HAB-forming |
| Cyanophyceae | *Synechococcus* sp. | WH 8102 | SN | 70-80 | 23 | No | oligotrophic |
| | *Synechococcus* sp. | WH 5701 | L1 + Si | 70-80 | 23 | Yes | coastal |
| | *Prochlorococcus marinus* | MIT9312 | Pro99 | 70 | 23 | Yes | high-light adapted |
| | *Prochlorococcus marinus* | NATL2A | Pro99 | 40 | 23 | Yes | low-light adapted |

**Supplementary Table 2. $eO_2^-$ irradiance curve parameters.** $eO_2^-$ production rates and irradiance data for each biological replicate were fit to a photosynthesis-irradiance model modified from [2] by Diaz et al. [1] (see Materials and Methods) * = data from Diaz et al. [1]

| Phytoplankton Strain | Biological Replicate | $P_D^{O2-}$ | $P_S^{O2-}$ | $P_m^{O2-}$ | α | β | $E_k^{O2-}$ | $R^2$ |
|---|---|---|---|---|---|---|---|---|
| | | amol cell$^{-1}$ h$^{-1}$ | | | $\frac{amol\ cell^{-1}h^{-1}}{\mu mol\ m^{-2}s^{-1}}$ | | µmol m$^{-2}$ s$^{-1}$ | |
| *Synechococcus* sp. WH 8102 | A | 21 | 56 | 68 | 0.053 | 0.002 | 1271 | 0.97 |
| | B | 35 | 3267 | 70 | 0.013 | 0.457 | 5222 | 0.83 |
| | C | 28 | 983 | 95 | 0.042 | 0.206 | 2261 | 1.00 |
| *Synechococcus* sp. WH 5701 | A | 16 | 1068 | 26 | 0.027 | 1.092 | 961 | 0.97 |
| | B | 14 | 209 | 20 | 0.024 | 0.285 | 837 | 0.88 |
| | C | 14 | 6 | 20 | 0.049 | 0.001 | 414 | 0.84 |
| *Prochlorococcus marinus* MIT9312 | A | 7 | 101 | 51 | 0.122 | 0.049 | 414 | 0.99 |
| | B | 2 | 1864 | 19 | 0.039 | 1.554 | 492 | 0.99 |
| | C | 7 | 368 | 52 | 0.108 | 0.272 | 483 | 0.99 |
| *Prochlorococcus marinus* NATL2A | A | 37 | 81 | 105 | 0.689 | 0.028 | 152 | 0.92 |
| | B | 16 | 48 | 50 | 0.287 | 0.030 | 175 | 0.91 |
| | C | 38 | 126 | 128 | 0.738 | 0.076 | 173 | 0.95 |
| *Emiliania huxleyi* CCMP 374 | A | 475 | 1107 | 1485 | 23.484 | 0.431 | 63 | 0.90 |
| | B | 162 | 575 | 641 | 11.282 | 0.499 | 57 | 0.89 |
| | C | 722 | 1186 | 1661 | 12.527 | 0.764 | 133 | 0.95 |
| *Emiliania huxleyi* CCMP 371 | A | 298 | 1099 | 1238 | 15.891 | 0.572 | 78 | 0.96 |
| | B | 247 | 704 | 891 | 22.836 | 0.411 | 39 | 0.98 |
| | C | 315 | 1194 | 1230 | 13.907 | 1.014 | 88 | 0.99 |
| *Ostreococcus tauri* OTH95 | A | 1 | 25286 | 163 | 0.930 | 52.801 | 175 | 1.00 |
| | B | 10 | 84543 | 191 | 1.169 | 200.841 | 163 | 0.97 |
| | C | 14 | 19863 | 225 | 1.401 | 47.809 | 160 | 0.97 |
| *Micromonas pusilla* CCMP1545 | A | 295 | 525 | 704 | 6.072 | 0.402 | 116 | 0.94 |
| | B | 235 | 305 | 505 | 6.958 | 0.182 | 73 | 0.97 |
| | C | 262 | 311 | 547 | 9.163 | 0.162 | 60 | 0.91 |
| *Dunaliella* sp. 15-1a | A | 4807 | 467192 | 7527 | 9.763 | 612.006 | 771 | 0.92 |
| | B | 5919 | 491118 | 9088 | 11.450 | 646.982 | 794 | 0.94 |
| | C | 4682 | 429379 | 6792 | 10.165 | 756.058 | 668 | 0.84 |
| *Symbiodinium* sp. CCMP 3364 | A | 0 | 186 | 180 | 35.457 | 0.168 | 5 | 0.97 |
| | B | 0 | 629 | 618 | 190.099 | 0.458 | 3 | 0.94 |
| | C | 0 | 571 | 563 | 143.464 | 0.287 | 4 | 0.90 |
| *Karenia brevis* ARC 5 | A | 105103 | 289013 | 270949 | 1644.497 | 342.555 | 165 | 0.89 |
| | B | 34694 | 263581 | 162532 | 569.618 | 179.279 | 285 | 0.95 |
| | C | 28387 | 88440426 | 143429 | 610.110 | 172241.168 | 235 | 0.93 |
| *Thalassiosira weissflogii* CCMP 1336 | A | 4298 | 1063152 | 21922 | 44.300 | 961.037 | 495 | 0.99 |
| | B | 1699 | 749719 | 21841 | 47.775 | 630.454 | 457 | 0.99 |
| | C | 3365 | 22667 | 14082 | 34.611 | 11.519 | 407 | 0.98 |
| *Thalassiosira pseudonanna* CCMP 1335 | A | 227 | 25891 | 1049 | 2.590 | 28.719 | 405 | 0.98 |
| | B | 44 | 64341 | 427 | 1.599 | 98.051 | 267 | 0.85 |
| | C | 37 | 15096 | 333 | 1.364 | 24.959 | 244 | 0.94 |
| *Thalassiosira oceanica* CCMP 1005* | A | 28 | 33 | 51 | 0.390 | 0.010 | 131 | 0.98 |
| | B | 17 | 42 | 58 | 0.340 | 0.002 | 168 | 1 |
| | C | 29 | 34 | 59 | 0.230 | 0.007 | 260 | 0.99 |
| *Phaeodactylum tricornutum* CCAP 1055/1 | A | 0 | 28 | 26 | 3.599 | 0.051 | 7 | 1 |
| | B | 0 | 66 | 62 | 13.292 | 0.159 | 5 | 1 |
| | C | 0 | 89 | 86 | 27.372 | 0.147 | 3 | 1 |
| *Aureococcus anophagefferens* CCMP 1984 | A | 19 | 9197 | 139 | 1.700 | 47.049 | 82 | 0.98 |
| | B | 41 | 205 | 178 | 3.358 | 0.438 | 53 | 0.99 |
| | C | 68 | 662 | 473 | 5.965 | 1.028 | 79 | 0.96 |

**Supplementary Table 3. Parameters for estimating $eO_2^-$ production rates under current and future conditions of the North Pacific Subtropical Gyre.** All data are obtained from the North Pacific gyre or another representative oligotrophic gyre. NPSG = North Pacific Subtropical Gyre, MLD = mixed layer depth

| Parameter (units) | Value | Notes | Source |
|---|---|---|---|
| Surface Par or $E_0$ Range (μmol $m^{-2}$ $s^{-1}$) | 324 – 613 | | Estimated from Figure 2 in Letelier et al. [3] |
| Surface PAR or $E_0$ Median (μmol $m^{-2}$ $s^{-1}$) | 481 | | Calculated with data from Letelier et al. [3] |
| Light attenuation coefficient (*k*; $m^{-1}$) | 0.04 | | Letelier et al.[3] |
| $5^{th}$ – $95^{th}$ percentile of the MLD (m) | 27 – 109 | | HOT-DOGS [9] |
| Median MLD (m) | 55 | | HOT-DOGS [9] |
| *Prochlorococcus* Abundance (cells $L^{-1}$) | $5.7 \times 10^7$ | Divided $10 \times 10^{12}$ $m^{-2}$ by listed depth integration (175 m) | Table 2 in Bjorkman et al. [10] |
| *Synechococcus* Abundance Range (cells $L^{-1}$) | $4.17 \times 10^5$ – $2.5 \times 10^6$ | Divided $0.5 \times 10^{11}$ $m^{-2}$ by shallowest (20 m) and deepest MLD (120 m) | HOT-DOGS [9] |
| Diatoms Abundance (cells $L^{-1}$) | $8.33 \times 10^2$ – $5 \times 10^3$ | Divided $1 \times 10^8$ $m^{-2}$ by shallowest (20 m) and deepest MLD (120 m) | Estimated from Figure 1 in Scharek et al. [11] |
| *Ostreococcus* Abundance (cells $L^{-1}$) | $10^3$ | | Figure 9 from Not et al. [12] |
| *Micromonas* Abundance (cells $L^{-1}$) | $7 \times 10^3$ – $21 \times 10^3$ | | Table VI in Furuya et al. [13] |
| Coccolithophore Abundance (cells $L^{-1}$) | $10^3$ | | Figure 3 in Cortés et al. [14] |

**Supplementary Table 4. Current and future ocean conditions estimated for the North Pacific Subtropical Gyre.** MLD = mixed layer depth, $E_0$ = surface irradiance or irradiance at 0 m, $E_z$ = irradiance at depth z, $P^{O_2^-}_{in\ situ}$ = production of $eO_2^-$ by *Prochlorococcus*

| | | $E_0$ = 324 µmol $m^{-2}$ $s^{-1}$ | | | $E_0$ = 481 µmol $m^{-2}$ $s^{-1}$ | | | $E_0$ = 613 µmol $m^{-2}$ $s^{-1}$ | | |
|---|---|---|---|---|---|---|---|---|---|---|
| **Ocean Conditions** | **MLD (m)** | $E_z$ (µmol $m^{-2}$ $s^{-1}$) | $P^{O_2^-}_{in\ situ}$ (nM $d^{-1}$) | $P^{O_2^-}_{in\ situ}$ (amol $cell^{-1}$ $h^{-1}$) | $E_z$ (µmol $m^{-2}$ $s^{-1}$) | $P^{O_2^-}_{in\ situ}$ (nM $d^{-1}$) | $P^{O_2^-}_{in\ situ}$ (amol $cell^{-1}$ $h^{-1}$) | $E_z$ (µmol $m^{-2}$ $s^{-1}$) | $P^{O_2^-}_{in\ situ}$ (nM $d^{-1}$) | $P^{O_2^-}_{in\ situ}$ (amol $cell^{-1}$ $h^{-1}$) |
| Future | 7 | 244.9 | 31 ± 15 | 23 ± 11 | 363.5 | 39 ± 19 | 29 ± 14 | 463.3 | 44 ± 21 | 32 ± 15 |
| Current | 27 | 110.0 | 20 ± 10 | 14 ± 7 | 163.3 | 25 ± 12 | 18 ± 9 | 208.2 | 28 ± 14 | 21 ± 10 |
| Future | 35 | 79.9 | 17 ± 8 | 12 ± 6 | 118.6 | 21 ±10 | 15 ± 7 | 151.2 | 24 ± 12 | 17 ± 9 |
| Current | 55 | 53.3 | 12 ± 6 | 9 ± 4 | 53.3 | 14 ± 7 | 10 ± 5 | 67.9 | 15 ± 8 | 11 ± 6 |
| Future | 89 | 9.2 | 9 ± 4 | 6 ± 3 | 13.7 | 9 ± 5 | 7 ± 3 | 17.4 | 10 ± 5 | 7 ± 4 |
| Current | 109 | 4.1 | 8 ± 4 | 6 ± 3 | 6.1 | 8 ± 4 | 6 ± 3 | 7.8 | 8 ± 4 | 6 ± 3 |

**Supplementary Table 5. Cell size and morphology data for surface area calculations of phytoplankton strains.** Radius was calculated as $\frac{width}{2}$. For prolate spheroids surface area calculations, $A = \frac{length}{2}$, $B = \frac{width}{2}$, and $m = \sqrt{1 - (\frac{B}{A})^2}$. NCMA = National Center for Marine Algae and Microbiota

| Genus species | Strain | Length (µm) | Width (µm) | Radius (µm) | Shape | Surface Area Formula | Surface Area (µm²) | Reference |
|---|---|---|---|---|---|---|---|---|
| *Emiliania huxleyi* | CCMP 374 | 6 | 5 | 2.5 | sphere | $4\pi r^2$ | 78.5 | NCMA |
| *Emiliania huxleyi* | CCMP 371 | 7 | 7 | 3.5 | sphere | $4\pi r^2$ | 153.9 | NCMA |
| *Ostreococcus tauri* | OTH95 | 1 | 0.7 | 0.35 | sphere | $4\pi r^2$ | 3.1 | Courties et al. [15] |
| *Micromonas pusilla* | CCMP 1545 | 3 | 2.5 | - | prolate spheroid | $2\pi B^2(1 + \frac{Asin^{-1}(m)}{mB})$ | 22.3 | NCMA |
| *Dunaliella* | 15-1a | 8.3 | 3.3 | - | prolate spheroid | $2\pi B^2(1 + \frac{Asin^{-1}(m)}{mB})$ | 71.6 | Microscopy |
| *Thalassiosira pseudonana* | CCMP 1335 | - | - | - | cylinder | $2\pi rh + 2\pi r^2$ | 51.0 | Kustka et al. [16] |
| *Thalassiosira weissflogii* | CCMP 1336 | - | - | - | cylinder | $2\pi rh + 2\pi r^2$ | 460.0 | Kustka et al. [16] |
| *Thalassiosira oceanica* | CCMP 1005 | - | - | - | cylinder | $2\pi rh + 2\pi r^2$ | 122.0 | Lommer et al. [17] |
| *Phaeodactylum tricornutum* | CCAP 1055/1 | 22 | 2.5 | - | pennate | - | 368.0 | Leblanc et al [18] |
| *Symbiodinium* sp. | CCMP 3364 | 13.5 | 11.5 | - | prolate spheroid | $2\pi B^2(1 + \frac{Asin^{-1}(m)}{mB})$ | 464.4 | Zhang et al. [19] |
| *Karenia brevis* | ARC 5 | 25 | 28 | 14 | sphere | $4\pi r^2$ | 2461.8 | Novoveská et al. [20] |
| *Aureococcus anophagefferens* | CCMP 1984 | - | - | 1.1 | sphere | $4\pi r^2$ | 15.2 | Sieburth and Johnson [21] |
| *Synechococcus* sp. | WH 8102 | 1.5 | 1.5 | 0.75 | sphere | $4\pi r^2$ | 7.1 | NCMA |
| *Synechococcus* sp. | WH 5701 | 1 | 1 | 0.5 | sphere | $4\pi r^2$ | 3.1 | NCMA |
| *Prochlorococcus marinus* | MIT9312 | 1.4 | 0.7 | - | prolate spheroid | $2\pi B^2(1 + \frac{Asin^{-1}(m)}{mB})$ | 2.6 | NCMA |
| *Prochlorococcus marinus* | NATL2A | 1.4 | 0.7 | - | prolate spheroid | $2\pi B^2(1 + \frac{Asin^{-1}(m)}{mB})$ | 2.6 | NCMA |
